# Real-time imaging reveals a role for macrophage protrusive motility in melanoma invasion

**DOI:** 10.1101/2024.09.25.614908

**Authors:** Gayathri Ramakrishnan, Veronika Miskolci, Miranda Hunter, Morgan A. Giese, Daniela Münch, Yiran Hou, Kevin W Eliceiri, Michael R. Lasarev, Richard M. White, Anna Huttenlocher

**Author notes:** Corresponding author: Anna Huttenlocher,.

## Abstract

Macrophages are primary cells of the innate immune system that mediate tumor progression. However, the motile behavior of macrophages and interactions with tumor cells are not well understood. Here, we exploit the optical transparency of larval zebrafish and perform real time imaging of macrophage-melanoma interactions. We found that macrophages are highly motile in the tumor microenvironment. Macrophages extend dynamic projections between tumor cells that precedes invasive melanoma migration. Modulating macrophage motility with a dominant inhibitory mutation in Rac2 inhibits recruitment to the tumor and impairs tumor invasion. However, a hyperactivating mutation in Rac2 does not affect macrophage recruitment but limits macrophage projections into the melanoma mass and reduces invasive melanoma cell migration. Taken together, these findings reveal a role for Rac2-mediated macrophage protrusive motility in melanoma invasion.

## Introduction

Tumor metastasis is a major cause of cancer-related deaths and yet the mechanisms that drive metastasis are incompletely understood (Dillekås et al., 2019). The role of innate immune cells in promoting metastasis is becoming increasingly evident (Kitamura et al., 2015; Gonzalez et al., 2018). Macrophages, primary cells of the innate immune system, regulate different steps of the metastatic cascade (Wyckoff et al., 2007; Goswami et al., 2005; Linde et al., 2018). For example, previous studies have shown that macrophages can guide tumor cells to the vasculature and mediate tumor cell movement across the endothelium (Arwert et al., 2018).

Macrophages are motile cells within interstitial tissues and display a mesenchymal surveillance mode of migration in human, mouse and larval zebrafish (Paterson & Lämmermann, 2022) (Barros-Becker et al., 2017). Rho GTPase signaling plays a critical role in regulating the dynamic actin cytoskeleton of motile macrophages. Depletion of Rac2, a hematopoietic specific Rac GTPase, in larval zebrafish impairs macrophage motility and recruitment to tissue damage (Rosowski et al., 2016). Rac2 depletion also results in decreased tumor burden and metastasis in mouse models (Joshi et al., 2014).

However, the relationship between macrophage migration in the tumor microenvironment (TME) and tumor invasion remains unclear. This gap is, in part, because of the difficulty in performing high resolution imaging of the TME *in situ*. Larval zebrafish are optically transparent and provide a powerful model to image dynamic interactions between tumor cells and macrophages in real-time *in vivo*. Melanoma is a highly aggressive type of skin cancer, and zebrafish melanoma recapitulates many of the properties of human melanoma tumors, including the ability to proliferate and invade (Heilmann et al., 2015). Indeed, a recent report used zebrafish to show that the anatomic position influences the transcriptional state of melanoma (Weiss et al., 2022). This progress highlights the ability of zebrafish melanoma models to uncover new mechanisms of disease progression. Macrophages have also been associated with melanoma dissemination in larval zebrafish and have been shown to transfer cytoplasm from macrophages to melanoma cells (Roh-Johnson et al., 2017).

Here we sought to determine how macrophages influence melanoma progression using live imaging. We used a previously established hindbrain injection model (Roh-Johnson et al., 2017) to evaluate the influence of macrophages on early melanoma invasion. Using real-time imaging, we showed the dynamic protrusive behavior and motility of macrophages within melanoma tumor cell masses. Depletion of macrophages abrogates tumor invasion. Expression of a dominant inhibitory mutation Rac2^D57N^ in macrophages led to inhibition of macrophage recruitment to the tumor and tumor invasion. Interestingly, hyperactivating mutations of Rac2 in macrophages affected macrophage protrusive motility in the tumor without altering recruitment to the tumor and impaired tumor invasion. Together, these findings suggest that macrophage protrusive motility in the TME impacts melanoma early invasion in zebrafish.

## Results and Discussion

### Macrophages preferentially accumulate near melanoma cells

To image interactions between innate immune cells and invasive melanoma cells, we used a previously established melanoma transplantation model in the larval zebrafish (Roh-Johnson et al.,2017). In this model, zebrafish melanoma cells injected into the hindbrain disseminate to distant sites by 3-4 days post injection (dpi) (Xue et al., 2023). We transplanted 15-20 zebrafish melanoma (ZMel1) cells derived from *mitfa-/-, mitfa:*BRAFV600E, *p53-/-* tumors (Heilmann et al.,2015) into the hindbrain ventricle of 2 days post fertilization (dpf) larvae (Figure 1A). To characterize the early events involved in tumor invasion, we imaged and tracked the dynamics of the transplanted tumor mass from 1 to 72 hours post injection (hpi). At 1 and 24 hpi, the tumor cells formed a clustered mass. However, by 48 hpi, tumor cells detached from the tumor mass and invaded into the surrounding tumor microenvironment (TME) (Figure 1B). The number of invaded tumor cells further increased by 72 hpi (Figure 1C). Quantification of the morphology of the tumor mass showed a significant reduction in the roundness of the mass by 48 hpi, suggesting remodeling of the tumor mass. (Figure 1D). This decrease in tumor roundness correlated with the onset of tumor cell invasion.

**Figure 1:**
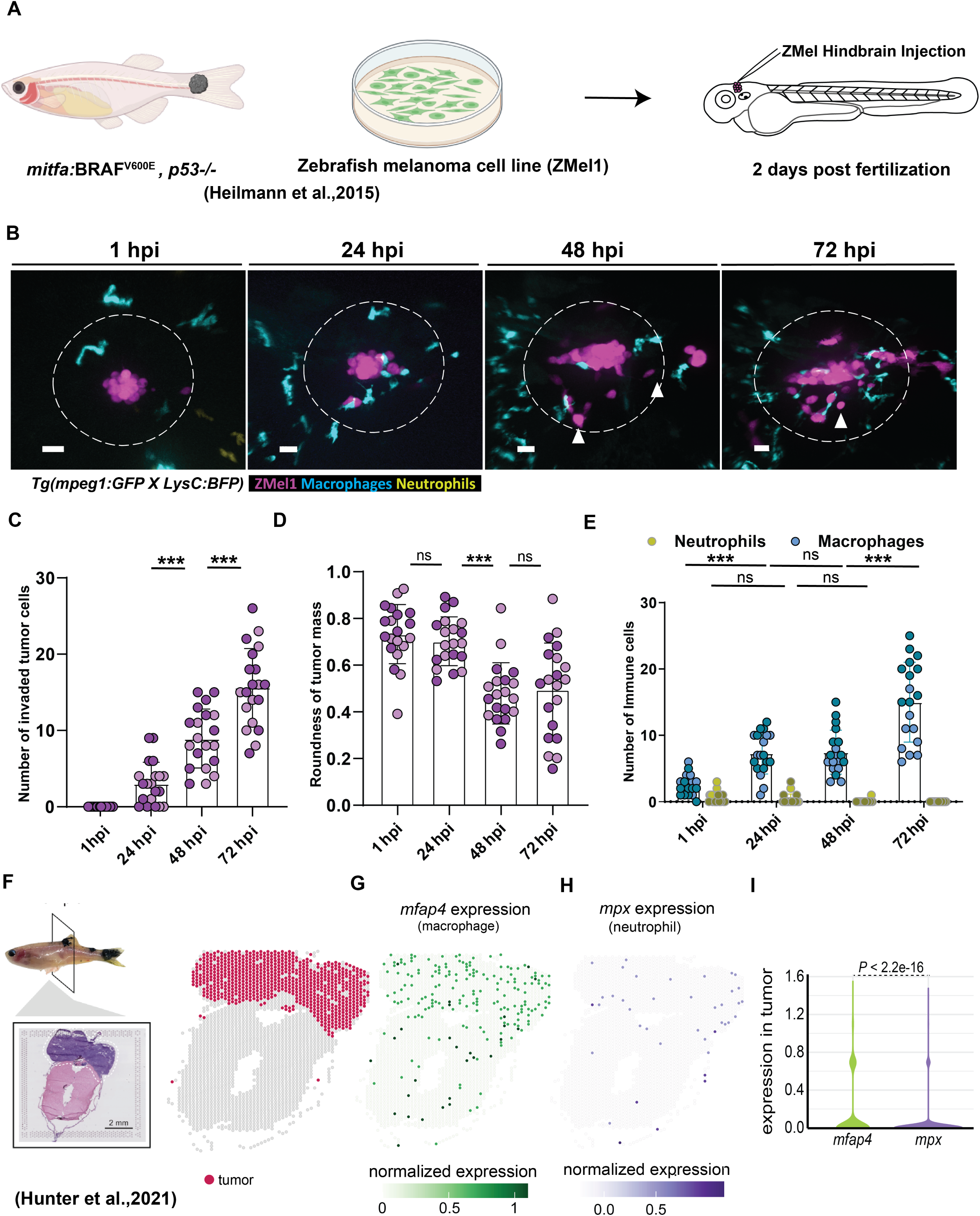
Macrophages preferentially accumulate near melanoma cells. **A.** Schematic showing the larval zebrafish melanoma model. ZMel cells were injected into 2 dpf zebrafish larvae *Tg(mpeg1:GFP, LysC:BFP)* and imaged from 1 to 72 hours post injection (hpi). **B.** Representative images of the melanoma tumor microenvironment (TME) from 1 hpi to 72 hpi. Arrows indicate invaded tumor cells. Dotted circle indicates region of immune cell quantification. **C.** Quantification of the number of invaded tumor cells over time. **D.** Quantification of the roundness of the tumor mass over time. **E.** Quantification of neutrophils and macrophages within 50 µm distance from the periphery of the tumor cluster. n=19 larvae from two independent experiments. Scale bar 20 µm. *p<0.05, **p<0.01, ***p<0.001. **F.** Image of zebrafish with *BRAF^V600E^*used for spatial transcriptomics in Hunter et al., 2021, Visium array spots color coded to indicate the tumor region. **G., H.** The expression of *mfap4* (macrophage marker) and *mpx* (neutrophil marker) projected over tissue space. **I.** Quantification of spatial gene expression in the tumor regions from 3 different tissue sections. Significance was calculated by Wilcoxon sum rank test.

Both neutrophils and macrophages can promote tumor invasion (Giese et al., 2019; Chen et al., 2019). We used zebrafish with labeled neutrophils *(LysC*:*BFP*) and macrophages (*mpeg1*:*GFP*) to characterize recruitment dynamics of innate immune cells over time (Figure 1B). At 1 hpi, there was recruitment of both neutrophils and macrophages. Over time, macrophages had an increase in recruitment, while neutrophil numbers decreased (Figure 1E). The increase in macrophage recruitment correlated with increased tumor cell invasion. Using time-lapse imaging, we found that neutrophils showed a transient presence around the tumor cells, while macrophages remained highly motile in the TME after transplantation of melanoma cells (Supplement Figure 1A, B).

To investigate if the limited recruitment of neutrophils is specific to the larval zebrafish, we assessed the expression of neutrophil and macrophage markers in a previously published spatial transcriptomics dataset of transgenic BRAF^V600E^-driven melanoma tumors in adult zebrafish (Figure 1F) (Hunter et al., 2021). We found a significantly lower expression of the neutrophil gene *mpx* compared to expression of macrophage-specific gene *mfap4* in the tumor area (Figure 1G-I), suggesting that macrophages are also enriched in adult zebrafish melanoma tumors.

### Macrophages mediate the invasive migration of melanoma cells

To characterize the role of macrophages in early melanoma invasion, we performed live imaging of macrophage-melanoma interactions following transplantation in the hindbrain. Long-term time-lapse imaging of the tumor mass was performed by acquiring images every 5 minutes for 12 hours, beginning at 24 hpi. Macrophages were highly motile around the tumor cells. Notably, macrophages migrated between tumor cells and extended elongated protrusions into the mass (Video 1). 3D rendering revealed macrophages infiltrating between tumor cells and single tumor cells migrating away from the tumor mass (Figure 2A). Over time, macrophages extended elongated protrusions between invading tumor cells before individual tumor cells separated from the mass (Figure 2B). Furthermore, macrophages extended longer protrusions while in contact with an invading tumor cell compared to contacting other cells in the tumor mass (Figure 2C). Using light sheet microscopy (LSM) and a membrane-labelled macrophage line (*mpeg:mCherry-CAAX*), we imaged interactions between tumor cells and macrophage projections. LSM was performed with image acquisition every minute at 48hpi for 2 hours. This imaging further demonstrated that macrophage protrusions extend between tumor cells during tumor invasion (Supplemental figure 1 C, video 2).

**Figure 2:**
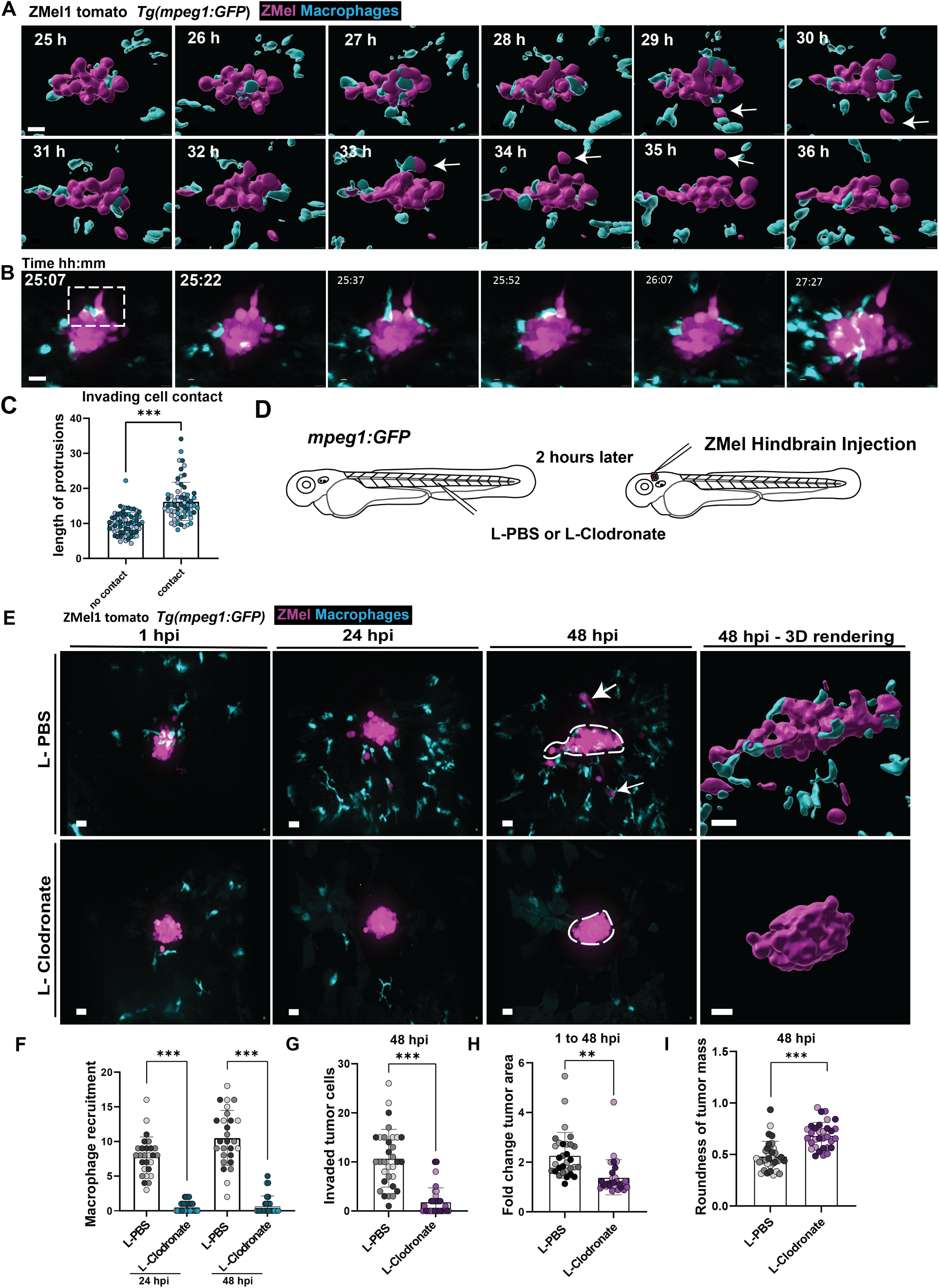
Macrophages mediate invasive migration of melanoma cells. **A.** Representative stills from time-lapse imaging of tumor-macrophage interactions between 24 and 36 hpi acquired every 5 minutes (Video 1). **B.** Representative stills of tumor-macrophage interactions that precede tumor invasion. **C.** Quantification of length of macrophage protrusions during macrophage contact with invading tumor cell compared to macrophages that do not contact the invading tumor cell. n=5 larvae; 27 contact cells and 25 non-contact cells from three independent experiments. Quantifications were made in single frame prior to each invasion event. **D.** Schematic of sequence of larvae injections. 2 dpf larvae were injected with L-PBS or L-Clodronate through the caudal vein followed by ZMel1 injection into the hindbrain ventricle 2 hours later**. E.** Representative images from time course imaging of L-PBS or L-Clodronate injected larvae from 1 hpi to 48 hpi. **F.** Quantification of macrophage recruitment at 24 and 48 hpi. **G.** Quantification of the number of invaded tumor cells at 48 hpi. **H.** Quantification of fold change in 3D tumor area from 1 hpi to 48 hpi. **I.** Quantification of roundness of tumor cell cluster at 48 hpi. n=30 larvae L-PBS, n=28 larvae L-Clodronate from three independent experiments. Scale Bar=20µm. *p<0.05, **p<0.01, ***p<0.001.

Previous studies showed that depletion of both macrophages and neutrophils in the zebrafish transplantation model limited tumor cell dissemination (Roh-Johnson et al., 2017). To identify a specific role for macrophages during tumor cell invasive migration, we depleted macrophages using clodronate liposomes. We injected either PBS or clodronate liposomes into the caudal vein of 2 dpf (*mpeg1:GFP)* larvae two hours prior to ZMel1 injections into the hindbrain (Figure 2D). We then imaged the same larvae every 24 hours (Figure 2E). Larvae that received clodronate had significantly reduced macrophages around the tumor cells at 24 hpi and 48 hpi (Figure 2F). When we measured invasion at 48 hpi, there was about a 10-fold reduction in the number of invaded tumor cells in macrophage depleted larvae compared to control (Figure 2G). The tumor area was also significantly reduced in the clodronate treated larvae (Figure 2H). Accordingly, the tumor mass was also significantly more rounded in clodronate treated larvae (Figure 2I). These data indicate that macrophages remodel the tumor mass and promote early invasion of melanoma cells.

### Rac2 signaling is required for macrophage recruitment to the TME and tumor invasion

To further study the effects of macrophage protrusions on melanoma invasion, we modulated Rac2 signaling which regulates F-actin assembly (Wheeler et al., 2006; Miskolci et al., 2016). We have previously shown Rac2 depletion results in defects in macrophage migration to a wound (Rosowski et al., 2016). We hypothesized that perturbing Rac2 signaling in macrophages would impair macrophage-mediated melanoma invasion. Indeed, we found that global deletion of Rac2 impaired melanoma invasion (Supplemental figure 2A, B). To modulate Rac2 signaling specifically in leukocytes, we expressed dominant inhibitory Rac2, Rac2^D57N^ (*Tg(coro1a:GFP-rac2_D57N)^psi92T^)*, under the coro1a promoter which is expressed predominantly in neutrophils and macrophages(Li et al., 2012). Since the neutrophil-specific expression of Rac2^D57N^ did not modulate tumor invasion (Supplemental figure 1A-C), the effects of Rac2^D57N^ using the coro1a promoter should be due to macrophage Rac signaling. Expression of Rac2D57N in macrophages restricted macrophages to the vasculature (Figure 3A) and impaired macrophage recruitment to a tail transection wound (Figure 3B-D). Macrophage expression of Rac2D57N also limited macrophage recruitment to the melanoma mass (Figure 3E, F). Accordingly, we found that this reduction in macrophage recruitment correlated with a decrease in tumor invasion and increased tumor roundness, suggesting impaired remodeling of the tumor mass with altered macrophage Rac2 signaling (Figure 3G,H). There was also a trend towards decreased tumor area (Figure 3I). These findings further support a central role for macrophages and macrophage Rac activation specifically in regulating macrophage motility and tumor invasion. Taken together, these results suggest that macrophage motility in the TME is required for early tumor invasion.

**Figure 3:**
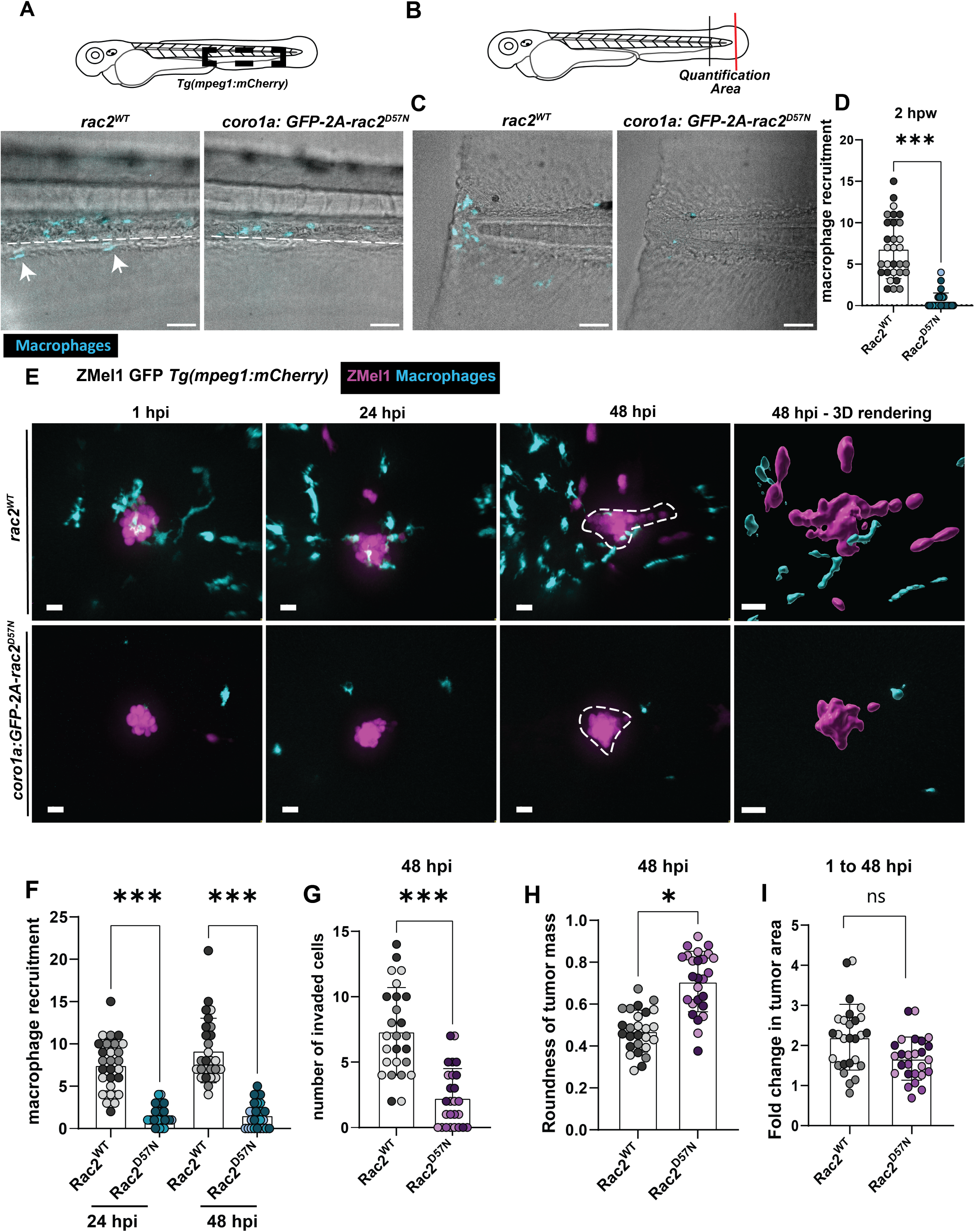
**Rac2 signaling in macrophages is required for macrophage recruitment to the TME and tumor invasion**. **A**, Representative images of macrophages in the caudal hematopoietic tissue of 3 dpf wild-type larvae or *Tg(coro1a:GFP- rac2_D57N)^psi92T^*larvae. **B.** Schematic of a 3 dpf larvae to highlight the tail transection region (red line) and the quantification area for macrophage recruitment (between red and black line). **C.** Representative images of tail fin of wild-type or Rac2^D57N^ larvae wounded and imaged 2 hours post wound. **D.** Quantification of macrophage recruitment to tail fin. n= 30 Wild-type, n= 27 Rac2^D57N^ from three independent replicates. **E.** Representative images from time course imaging of wild-type or Rac2^D57N^ larvae injected with ZMel GFP cells from 1hpi to 48hpi. **F.** Quantification of macrophage recruitment to tumor at 24 and 48 hpi. **G.** Quantification of number of invaded tumor cells at 48 hpi. **H.** Quantification of tumor roundness at 48 hpi. **I.** Quantification of fold change in tumor area from 1 to 48hpi. n=26 wild-type, n=26 Rac2^D57N^ from three independent replicates. Scale Bar = 20µm. *p<0.05, **p<0.01, ***p<0.001, n.s- not significant

### Hyperactivation of Rac2 alters macrophage morphology and motility in the TME

We next sought to modulate macrophage specific Rac2 signaling by hyperactivating Rac2 signaling as we hypothesized dysregulated Rac2 may alter macrophage motility in the TME and tumor invasion. To do this, we expressed an activating Rac2^E62K^ mutation specifically in macrophages that has been reported in patients with immune dysregulation (Hsu et al., 2019). Previous studies have demonstrated that this disease- associated mutation results in an increase in Rac2 activation and hyperactivation of macrophages (Mishra et al., 2023). To determine if Rac2^E62K^ affects macrophage motility, we first assessed macrophage migration to a wound. We performed caudal fin tail transection at 3 dpf on larvae expressing *mpeg1:mCherry-2A-rac2^WT^*or *mpeg1:mCherry-2A-rac2^E62K^* (Supplemental figure 3D). We found that at 2 hours post wound (hpw) there was no difference in macrophage recruitment to a wound or change in the morphology of macrophages (Supplemental figure 3E, F).

We then assessed the effect of Rac2^E62K^ expression on macrophage recruitment to melanoma cells. Macrophage recruitment was not altered in macrophages expressing Rac2^E62K^ (Figure 4A, B). While, the roundness of the tumor mass and fold change of tumor area were not changed, we did observe a significant reduction in the number of invaded tumor cells in larvae expressing Rac2^E62K^ in macrophages (Figure 4C-E). Furthermore, macrophages exhibited a more rounded morphology with fewer and shorter protrusions near tumor cells in the Rac2^E62K^ mutants (Figure 4F-I). These data indicate that expression of Rac2^E62K^ mutation does not alter macrophage recruitment but changes macrophage protrusive morphology that could influence the interactions between macrophages and tumor cells. Interestingly, this effect was important in the tumor but not at the wound, suggesting distinct functions for macrophage Rac2 signaling at the wound and TME.

**Figure 4:**
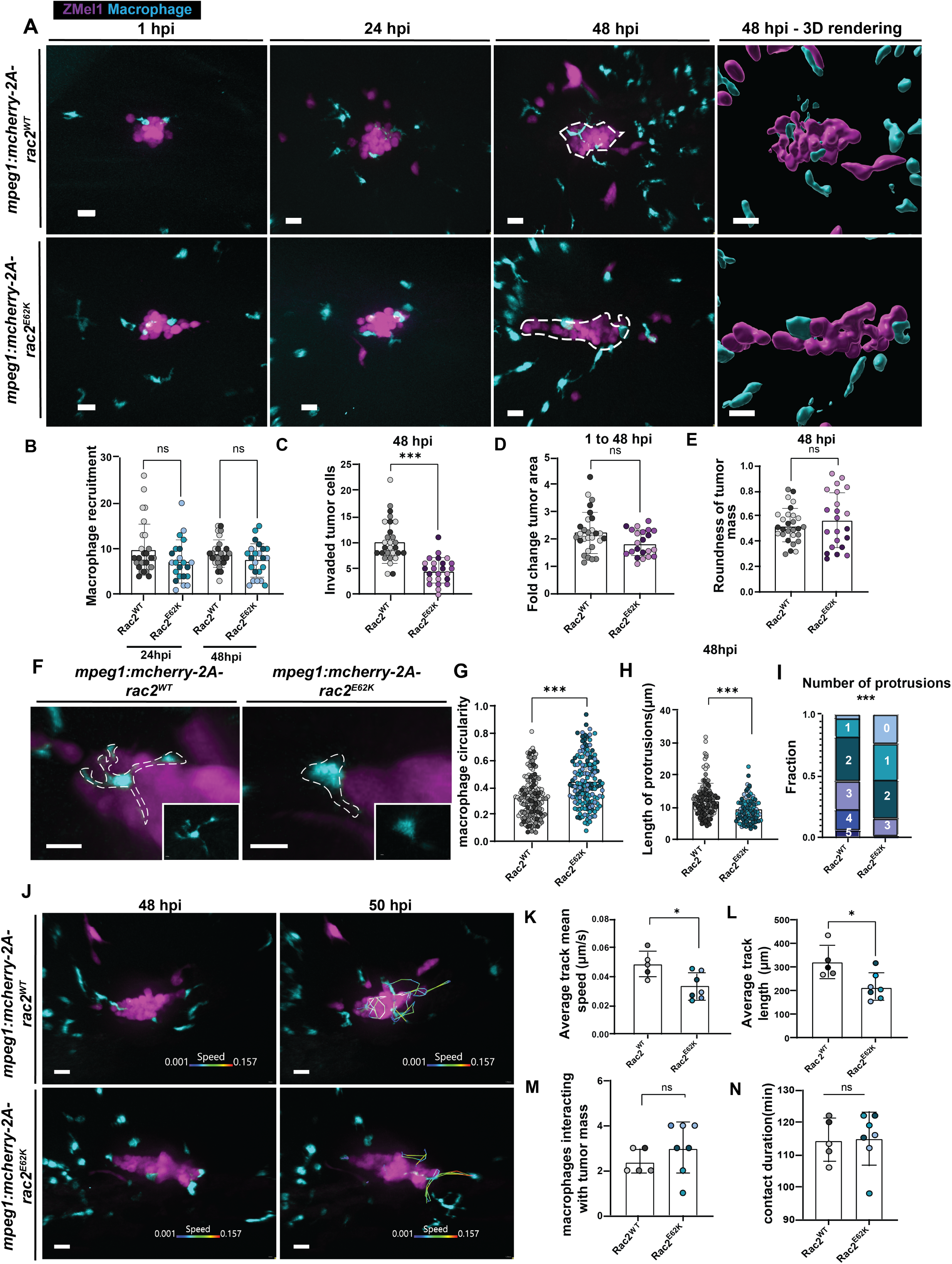
Hyperactivation of Rac2 alters macrophage morphology and motility in the TME. **A.** Representative images from time course of larvae expressing *mpeg1:mCherry-2A-rac2^WT^*, *mpeg1:mCherry-2A-rac2^E62K^* injected with ZMel1 cells at 2 dpf. **B.** Quantification of macrophage recruitment to tumor cells at 24 and 48 hpi. **C.** Quantification of the number of invaded tumor cells at 48 hpi. **D.** Quantification of roundness of tumor cell cluster at 48 hpi. **E.** Quantification of fold change in tumor area from 1 hpi to 48 hpi. Rac2^WT^ (n=28 larvae), Rac2^E62K^ (n=23 larvae) from three independent experiments. **F.** Zoom-in images of macrophages highlighting protrusions into the tumor mass in Rac2^WT^ and Rac2^E62K^-expressing larvae. **G., H., I.,** Quantification of circularity, length of protrusions and number of protrusions for macrophages at 48 hpi. Number of cells- Rac2^WT^= 161 cells, Rac2^E62K^= 145 cells. **J.** Representative images from time-lapse imaging of tumor-macrophage interactions at 48 hpi (Video 3). Tracks indicate macrophage motility around the tumor cells color-coded by instantaneous speed. **K.** Quantification of average track mean speed of macrophages from each larva. **L.** Quantification of average track length from macrophages in each larva. **M**. Quantification of the number of macrophages that interact with the tumor mass in the time course. **N.** Quantification of duration of contact of macrophages with the tumor mass in the time course. Rac2^WT^ (n=5 larvae; 12 cells), Rac2^E62K^ (n=7 larvae, 21 cells) from three independent experiments. Scale Bar=20 µm. *p<0.05, **p<0.01, ***p<0.001

To determine if the dynamic activity of macrophages in the TME was altered with Rac2^E62K^, we performed time-lapse imaging at 48 hpi. Macrophages expressing Rac2^E62K^ exhibited a slower migration speed and migrated a shorter distance in the 2- hour imaging window around the tumor cells (Figure 4J-L; Video 3). Time-lapse movies also revealed that mutant macrophages extended shorter and fewer protrusions into the tumor mass than wild-type macrophages that extended sustained elongated protrusions into the tumor mass (Video 3). However, we did not observe a significant difference in the number of macrophages that made contact or the contact duration with the tumor mass between wild-type and Rac2^E62K^ macrophages (Figure 4M,N). Our findings suggest that changes in macrophage behavior and the defects in protrusion formation with expression of Rac2^E62K^ is associated with defects specifically in tumor invasion. Taken together, these results suggest that regulated Rac2 signaling is necessary for the formation of elongated macrophage protrusions that extend into the tumor mass and promote tumor invasion.

Here, we take advantage of the live imaging capabilities of larval zebrafish to visualize the migratory behavior of macrophages in response to tumor cells in an *in vivo* environment. We found that melanoma invasion correlates with the presence of macrophages but not neutrophils in the TME. Indeed, depletion of macrophages, but not neutrophils, blocked tumor remodeling and invasion. Our results are consistent with findings in breast cancer that show early infiltration of macrophages into the tumor disrupts E-cadherin junctions and modulates tumor dissemination (Linde et al., 2017). In Drosophila embryos macrophages also penetrate developing tissues specifically during cell division, when there is a release of cell-cell adhesion (Akhmanova et al., 2022). These studies suggest that macrophages have mechanisms to migrate between contacting cells and this behavior may weaken cell-cell adhesion and facilitate tumor infiltration. Morphometric analyses in our study revealed that the melanoma mass became elongated and remodeled over time in response to macrophage infiltration. Macrophages likely mediate single-cell tumor invasion by disrupting cadherin-mediated cell-cell contacts, but other mechanisms are also possible. Secreted signaling factors, cytoplasmic transfer (Roh-Johnson et al., 2017) or tunneling nanotubes between macrophages and tumor cells may also play a role (Hanna et al., 2019).

Our findings demonstrate that macrophages that express Rac2^E62K^ have altered motility and projections into tumors but not wounds. The defects in protrusion formation in macrophages in Rac2^E62K^ could be due to impaired actin cycling (Hsu et al., 2019). It is possible that other mechanisms contribute to the effects of macrophage Rac2^E62K^ on tumor cell behavior. For example, macrophages expressing Rac2^E62K^ have recently been shown to exhibit an increased ability to phagocytose tumor cells *in vitro* (Mishra et al., 2023). Although, we did not observe hyperphagocytosis by Rac2^E62K^ macrophages by live imaging, more sensitive assays may be required to further evaluate this phenotype. Taken together, our findings raise the interesting possibility that motile macrophages in the TME provide a tumor invasion promoting role.

Previous studies using mouse models have shown that macrophages and tumor cells exhibit a leader-follower type of interaction where macrophages lead tumor invasion (Wyckoff et al., 2007)(Roussos et al., 2011). By utilizing high resolution live imaging methods, our study reveals a role for macrophage protrusions in promoting tumor invasion by infiltrating into the tumor and promoting tumor cell invasion from the tumor mass. In in vitro spheroid models, both amoeboid and mesenchymal modes of macrophage migration have been implicated in infiltrating the tumor (Guiet et al., 2011). Our study in a zebrafish model reveals elongated protrusions of macrophages associated with a mesenchymal motility mode during tumor infiltration, that is mediated by Rac2 signaling. Our study adds to the growing body of evidence for the tumor-promoting role of macrophages and provides new insights into the potential mechanism of these effects. This model could be further exploited to study the signaling mechanisms by which macrophages promote invasive behavior of tumor cells. Future studies should also further evaluate the relationship between macrophage polarization phenotype and motility in the TME and tumor invasion.

## Acknowledgements

We thank Alexandra Fister and Julie Rindy for technical assistance and members of the Huttenlocher lab for helpful discussions. We also thank Joe Li, Liz Haynes, and Kurt Weiss for assistance with light sheet microscopy. We would also like to thank Tatjana Piotrowski for sharing the Coro1A- Rac2D57N line.

## Data Availability

All relevant data are included in the manuscript.

## Author Contributions

G.R. and A.H. conceived and designed experiments. G.R., V.M., M.A.G. performed experiments. M.H. performed bioinformatics analysis. M.R.L. performed statistical analysis. G.R., V.M., M.A.G., M.H., R.M. and A.H. analyzed and interpreted the data.

G.R. and A.H. wrote the paper.

## Methods

### Zebrafish Lines and Maintenance

Animal care and use protocol M005405-A02 was approved by the University of Wisconsin-Madison College of Agricultural and Life Sciences (CALS) Animal Care and Use Committee. Adult zebrafish and larvae were maintained as previously described. Larvae were maintained in E3 containing 0.2 mM N-phenylthiourea beginning 24 hours post fertilization (PTU, Sigma Aldrich) to prevent pigment formation in the larvae. Larvae were anesthetized in E3 water containing 0.2 mg/mL Tricaine (ethyl 3-aminobenzoate, Sigma) prior to experimental procedures. The following fish lines were used for the study:

|  |  |
| --- | --- |
| *Tg(mpeg1:GFP)* | (Ellett et al., 2011) |
| *Tg(mpeg1:mCherry)* | (Ellett et al., 2011) |
| *Tg(mpeg:mCherry-CAAX)* | (Bojarczuk et al., 2016) |
| *Tg(mpx:mCherry-2A-rac2<sup>D57N</sup>)* | (Deng et al., 2011) |
| *Tg(coro1a:GFP-rac2_D57N)<sup>psi92T</sup>* | This paper |
| *Tg(mpeg1:mCherry-2A-rac2<sup>WT</sup>)* | (Rosowski et al., 2016) |
| *Tg(mpeg1:mCherry-2A-rac2<sup>E62K</sup>)* | This paper |
| *Tg(LysC:BFP)* | (Rosowski et al., 2018) |
| *Tg(mpeg1:GFP- histone2b)* | (Miskolci et al., 2019) |

### Morpholino injection

We used a previously published morpholino against *arnt-1* 5′-GGATTAGCTGATGTC ATGTCCGACA-3′ (Prasch et al., 2006) obtained from Gene Tools and diluted to a concentration of 500 µM. 3nL volume was injected at one-cell stage in zebrafish embryos. Non- targeting morpholino was used as a control.

### Generation of *Tg(mpeg1:mCherry-2A-rac2^E62K^)* transgenic line

*mCherry-2A-rac2^WT^* fragment was cut from existing construct (Deng et al., 2011) and inserted into pCRII TOPO vector via BamHI and NotI restriction sites (New England Biolabs, NEB). E62K (Glutamic acid to Lysine) mutation was generated by introducing a G to A point mutation at nucleotide 184 in the zebrafish Rac2 gene by QuikChange II Site-Directed Mutagenesis (Agilent kit) using primers 5’- cagtctgtcataatctttctgtccggctgtatccc-3’ and 5’-gggatacagccggacagaaagattatgacagactg-3’ (anti-sense).

Primers were designed using Agilent primer design tool at https://www.agilent.com/store/primerDesignProgram.jsp. The point mutation was confirmed by sequencing. *mCherry-2A-rac2^E62K^* fragment was cut and inserted into Tol2-*mpeg1* backbone vector (Ellett et al., 2011) using BamHI and NotI restriction sites. 3 nL of mixture containing 25 ng/μL plasmid DNA, 35 ng/μL Transposase mRNA and 0.5X CutSmart Buffer (NEB) was microinjected into the cytoplasm of one-cell stage embryos. mRNA was *in vitro* transcribed using mMESSAGE mMACHINE SP6 (Ambion kit) from a pCS2-transposase vector. Larvae were screened for mCherry expression in macrophages at 3 dpf and grown up to establish a stable line. Founders were identified by outcrossing to wild-type AB fish.

### Generation of the *Tg(coro1a:GFP-rac2D57N)^psi92Tg^* transgenic line

A DNA sequence 7.03kb upstream of the zebrafish *coro1a* translation start site was amplified by PCR as previously described (Li et al., 2012) and cloned into the tol2kit p5E-MCS vector (Kwan et al., 2007) using Gibson assembly (New England Biolabs). A dominant-negative variant of zebrafish *rac2* harboring a point mutation (D57N, (Deng et al., 2011)) was cloned from a gBlock HiFi Gene Fragment (Integrated DNA Technologies) containing coding DNA sequences for GFP and rac2_D57N, separated by a viral P2A peptide (GFP-P2A-rac2_D57N). The GFP-P2A-rac2_D57N fragment was amplified using primers with Gibson overhangs.

(5’-gtatcgataagcttgatatcgaattcaccatggtgtccaagggcgagga-3’ and 5’gtggatcccccgggctgcaggaattttagagcatcacgcagccct-3’) and cloned into the tol2kit pME-MCS vector (Kwan et al., 2007)linearized with EcoRI using Gibson assembly. The final expression vector was assembled by combining p5E-coro1a, pME-GFP-P2A- rac2_D57N, p3E-polyA (Kwan et al., 2007) and pToneDest (Shin et al., 2016) using Gateway LR Clonase II (Thermo Fisher Scientific). Tol1 mRNA was synthesized from pToneTP vector (Shin et al., 2016) using mMessage T7 kit (Thermo Fisher Scientific). Both pToneDest and pToneTP were gifts from Nathan Lawson (Addgene plasmids #67691 and #67692).

Zebrafish one-cell stage embryos were injected with Tol1 mRNA (40ng/ul) and expression vector (15ng/uL) and screened for GFP fluorescence at 2 dpf. Founders were identified based on GFP expression in immune cells and outcrossed to wildtype fish to generate stable F1 generations. Experiments were performed on larvae obtained from the F3 generation.

### Zebrafish melanoma cell line maintenance

We utilized ZMel1-GFP or ZMel1-tomato cell lines generated from a primary zebrafish melanoma model expressing BRAF V600E in a p53-/- background (Heilmann et al., 2015)(Campbell et al., 2021). ZMel1 cells were cultured in 10% FBS-DMEM +1% Glutamax +1% penicillin-streptomycin on 10 µg/mL fibronectin coated plates in a sterile 28°C incubator. To harvest, cells were washed with sterile PBS, harvested with trypsin and then counted with a hemocytometer.

### ZMel1 hindbrain injection

We modified a previously established hindbrain injection model of melanoma cells (Roh- Johnson et al., 2017). Briefly, cells were harvested, washed in PBS, then resuspended in HBSS media at a concentration of 80 million cells/mL. Cells were then loaded into thin-walled glass capillary injection needles. The needle was then calibrated to inject 1 nL volume (15-20 cells). Anesthetized larvae were then placed on 3% Agarose plate made with E3 and microinjected with ZMel1 cells, with time range set to “millisecond” and pressure set to ∼15 psi on the microinjector.

### Macrophage depletion

Macrophage depletion was performed by injection of clodronate or PBS liposomes (www.clodronateliposomes.com). At 2 dpf, anesthetized (*Tg(mpeg1:GFP)*) larvae were injected intravenously via the posterior caudal vein with 1 nL liposome-encapsulated clodronate (Rosowski et al., 2016) or PBS. After 2 hours, liposomes injected larvae were injected with ZMel1 cells in the hind brain ventricle.

### Zebrafish tail wounding

Dechorionated larvae 3 dpf larvae were tricaine - anesthetized and transferred to milk- coated 35 mm petri dishes. Tail wounding was performed in 0.2 mg/mL tricaine/E3 solution by transecting the caudal tail fin distal to the notochord using a #10 scalpel blade. Larvae were then allowed to recover by placing in E3 without tricaine at 28.5°C until imaging.

### Image acquisition

#### Invasion assay

To assess invasion at the injection site, larvae were anesthetized and mounted in a zWEDGI device (Huemer et al., 2017)such that the hindbrain was fully visible. Z-series images (3.45 µm) of the hindbrain were acquired on a spinning disk confocal microscope (CSU-X; Yokogawa) with a confocal scan head on a Zeiss Observer Z.1 inverted microscope, Plan-Apochromat NA 0.8/20x objective, and a Photometrics Evolve EMCCD camera. Between imaging session larvae were kept in E3 with PTU in individual 24-well plates. Individual larvae were followed throughout the time course. Images of larvae in the figures represent a 3D rendering of the images generated on Imaris 10.0.

#### Time-lapse imaging

To characterize macrophage behavior around tumor cells, time-lapse imaging was performed at 24 hpi (Figure 3) or 48 hpi (Figure 4). Larvae were loaded on a zWEDGI device and embedded with 1.5% low melting agarose (Fisher Scientific). Agarose was allowed to solidify and Tricaine/E3-PTU solution was then added to the device. All time- lapse movies were acquired on a spinning disk confocal microscope (CSU-X; Yokogawa) with a confocal scan head on a Zeiss Observer Z.1 inverted microscope, Plan-Apochromat NA 0.8/20x objective, and a Photometrics Evolve EMCCD camera. To characterize interactions between macrophages and tumor cells prior to invasion (Figure 2), 12-hour time-lapse imaging acquiring every 5 minutes starting at 24 hpi. To quantify macrophage motility parameters, time-lapse movies were performed at 24 and 48 hpi for 2 hours acquiring every 3 minutes. Multi time-lapse images were acquired using Zen software (Zeiss).

#### Light sheet Microscopy

To prepare the sample for imaging, FEP tube with inner diameter at 0.8 mm and outer diameter at 1.2 mm were cleaned as described(Weber et al., 2021). 1.2% and 2% low melting point (LMP) agarose stocks were preheated to 75°C then cooled down to 37°C. Tricaine methanesulfonate (MS222) stock solution was added to 1.2% LMP agarose to reach a final concentration of 200 mg/L. Zebrafish embryos were anesthetized in 1×E3 with MS222 at 200 mg/L, and mounted with 1.2% LMP agarose inside cleaned FEP tubes. The end of the tubes were sealed with 2% LMP agarose plugs. FEP tubes with mounted zebrafish embryos were kept in 1×E3 with MS222 before imaging. Sample tubes were loaded onto the sample holder and submerged into the sample chamber containing 1×E3 with MS222 heated up to 28°. Samples were imaged with 32x objective. Images were acquired every minute for 2 hours. Z-stack data sets were acquired on a PCO Panda 4.2M camera system running at 40 frames per second (FPS). Each image is 2040 x 2048 pixels at 16 bit pixel depth. Image acquisition is synchronized with a hardware transistor transistor logic (TTL) triggering signal between the camera, stages, and laser engine system. Stage xyz motion is controlled by a PI 884-C controller and M110 stages. The laser engine is a Toptica CLE capable of running at 640 nm 50 mW, 561 nm 40 mW, 488 nm 50 mW, and 405 nm 40 mW.

### Quantifications

#### Invaded ZMel1 cells and immune cell recruitment to ZMel1

Images acquired 1, 24 and 48 hpi were 3D rendered on Imaris 10.0. Invaded tumor cells were quantified as cells that were fully separated from the injected tumor cell mass at 48 hpi. For neutrophil and macrophage recruitment, larvae expressing *Tg(mpeg1:GFP) x Tg(lysC:BFP)* were used and cells present at 50 µm from the injected tumor mass were quantified.

#### ZMel1 area

Fold change in tumor area was quantified on Imaris 10.0. Surface function was used to create 3D surfaces on tumor region using the fluorescent signal with background subtraction. Total area of the tumor cells from each larva were obtained. Fold change in area from 1 to 48 hpi was then calculated and plotted.

#### ZMel1 cell mass and macrophage morphology

Morphology analysis was performed in Fiji(Weber et al., 2021). To calculate tumor roundness, images were z-projected, and tumor fluorescent signal was used to threshold. Using the wand tool, only the tumor mass region was selected and roundness (roundness=4×area/π×major axis^2^) was measured. Similarly, to calculate circularity of macrophages which indicates cell spread, images were first z projected. Macrophage fluorescent signal was then used to threshold and wand tool was used to select macrophages within 30µm distance from the tumor mass to measure circularity (circularity=4 π(area/perimeter^2^). To quantify macrophage protrusions number and length, images were 3D rendered on Imaris 10.0 software. Number of protrusions were measured by counting the number of extensions from the cell body. Length of protrusions were calculated by manually drawing a line from the cell body to the end of the extensions.

#### Cell Tracking

To track macrophage motility parameters, time-lapse images were loaded on Imaris 10.0. Spot function was used to track macrophages where spot size was defined as 6.5 µm diameter. Tracks were synthesized by manually tracking the cells through the duration of the time-lapse. Measurements such as track mean speed and track length were then obtained and plotted as average for each larva. Macrophages that interact with the tumor cluster were tracked for *mpeg1:mCherry-2A-rac2^E62K^* experiments.

Macrophages within 50 µm distance was tracked for *arnt-1* MO experiments. The number of tracks infiltrating the ZMel1 cell mass was measured by using surface-spot interactions statistics where tumor cells mass is surface rendered, and macrophages are tracked using spots. Track is considered infiltrating the tumor cell mass when it has a negative distance of the track from the tumor surface.

#### Macrophage recruitment to wound

Macrophage recruitment was quantified by counting cells in the caudal fin tissue area distal to the caudal vein loop using the fluorescent signal.

### Re-analysis of zebrafish melanoma spatial transcriptomics data

Zebrafish melanoma spatially resolved transcriptomics data from [PMID: 34725363] was analyzed as previously described (Hunter et al., 2021). All analysis was done in R (version 4.3.1) using Seurat [PMID: 37231261] (versions 4.4.0 and 5.3.1) (Hao et al., 2024). Briefly, raw counts were normalized using SCTransform [PMID: 31870423] (Hafemeister & Satija, 2019). The three biological samples were integrated using the Seurat workflow for integration of SCTransform-normalized data. PCA and UMAP dimensionality reduction were calculated using default parameters and cell type annotations were done using expression of typical marker genes. Spatial gene expression was plotted using the Seurat function SpatialPlot applied to the normalized counts. The Wilcoxon rank sum test was used to calculate differences in tumor expression between genes. The complete dataset is available from GEO (accession number GSE159709).

### Statistical analysis

All graphs were plotted on Graphpad (Prism). Different colored data points represent replicates, defined as clutches of larvae spawned on different days. All statistical analyses were performed in collaboration with a statistician. Count phenomenon (number of protrusions, invasion, macrophages/neutrophils recruited, infiltrating tracks) were analyzed using Poisson regression with cluster-robust standard errors to account for subsampling (larvae from the same clutch, macrophages within larvae). If count data exhibited greater variation than predicted from a Poisson distribution, then quasi- likelihood was used to fit the (Poisson) regression models with standard errors inflated by an estimated multiplicative factor (>1) of dispersion. Continuous outcomes (circularity, area, roundness, motility.) were analyzed using linear models with cluster- robust standard errors. Responses such as area fold change were log-transformed prior to fitting the linear model. Experimental results for the time course experiment were fit using mixed-effect models (for the Poisson or Gaussian family, depending on whether the response was a discrete count or continuous) to account for repeated measures over time on the same unit. Half-normal plots with a simulated envelope and plots of residuals vs fitted values were used to check adequacy of fitted models.

---

**Supplemental figure 1:** Macrophages accumulate and extend protrusions into the tumor mass correlated with tumor invasion. **A.** Representative stills from time-lapse imaging of ZMel1 transplanted larvae from 1 hpi to 6 hpi. **B.** Quantification of neutrophils and macrophages every hour from 1-6 hpi indicated by average values at each time point. n=4 larvae. C. Representative z-projected stills from LSM imaging of tumor-macrophage interactions at 48hpi imaged every minute. Lines indicate macrophage protrusions and invading tumor cell (n=4).

**Supplemental figure 2:** Depletion of Rac2 inhibits tumor invasion. **A.** Representative stills from time course imaging of ZMel1 cells after transplantation in *rac2^+/+^*or *rac2^-/-^*. **B.** Quantification of number of invaded tumor cells in *rac2^+/+^* (n=18 larvae), *rac2^-/-^* (n=23 larvae) and *rac2+/-* (n=37 larvae) from three independent replicates. *p<0.05, **p<0.01.Scale Bar= 20 µm

**Supplemental figure 3:** Effects of Rac2 on tumor invasion and macrophage motility at wounds. **C.** Representative stills from time course images of wild-type larvae or larvae expressing *mpx:mCherry-2A-rac2^D57N^* injected with ZMel1 cells at 2 dpf. **D.** Quantification of number of invaded tumor cells at 48 hpi. **E.** Quantification of macrophage recruitment at 24 and 48 hpi. wild-type (n= 18 larvae), *mpx:mCherry-2A- rac2^D57N^* (n=20 larvae). Scale Bar= 20 µm **C.** Representative stills of *mpeg1:mCherry- 2A-rac2^WT^*or *mpeg1:mCherry-2A-rac2^E62K^* expressing larvae at 2 hours post wound (hpw), tail transected at 3 dpf. **D.** Quantification of macrophage recruitment to wound at 2 hpw. E. Quantification of circularity of macrophages at the wound. *mpeg1:mCherry- 2A-rac2^WT^* (n=21 larvae, 219 cells), *mpeg1:mCherry-2A-rac2^E62K^* (n= 22 larvae, 207 cells) from two independent experiments. *p<0.05, **p<0.01.

**Video 1: Imaging macrophage motility in the tumor microenvironment**. Time-lapse movie acquired every 5 minutes for 12 hours from 24 hpi of ZMel1 cells in *mpeg1:GFP* line. n= 5 larvae. Video was generated on Imaris 10.0. frame rate- 5fps. Scale = 10 µm.

**Video 2: LSM imaging of macrophage motility and tumor invasion**. Time-lapse movie acquired every minute for 2 hours at 48hpi of ZMel1 cells in *mpeg1:mcherry-caax* line. Video was generated on Imaris 10.0. frame rate – 5fps. Scale = 10 µm.

**Video 3: Effect of Rac2^E62K^ on macrophage motility in the TME**. Larvae expressing *mpeg1:mCherry-2A-rac2^WT^* or *mpeg1:mCherry-2A-rac2^E62K^*were injected with ZMel1 cells at 2 dpf and time-lapse imaging was performed at 48 hpi. Images were acquired every 3 minutes for 2 hours. Macrophages that interact with the tumor mass were tracked and color coded by instantaneous speed of macrophages. Videos were generated on Imaris 10.0 and processed on premier pro. Rac2^WT^ (n=5 larvae), Rac2^E62K^ (n=7 larvae) from three independent replicates. frame rate- 5fps. Scale bar = 10µm.

## References

1. Arwert, E. N., Harney, A. S., Entenberg, D., Wang, Y., Sahai, E., Pollard, J. W., & Condeelis, J. S. (2018). A Unidirectional Transition from Migratory to Perivascular Macrophage Is Required for Tumor Cell Intravasation. Cell Reports, 23(5), 1239– 1248. 10.1016/j.celrep.2018.04.007

2. Barros-Becker, F., Lam, P.-Y., Fisher, R., & Huttenlocher, A. (2017). Live imaging reveals distinct modes of neutrophil and macrophage migration within interstitial tissues. Journal of Cell Science, 130(22), 3801–3808. 10.1242/jcs.206128

3. Bojarczuk, A., Miller, K. A., Hotham, R., Lewis, A., Ogryzko, N. V., Kamuyango, A. A., Frost, H., Gibson, R. H., Stillman, E., May, R. C., Renshaw, S. A., & Johnston, S. A. (2016). Cryptococcus neoformans Intracellular Proliferation and Capsule Size Determines Early Macrophage Control of Infection. Scientific Reports, 6(1), 21489. 10.1038/srep21489

4. Campbell, N. R., Rao, A., Hunter, M. V., Sznurkowska, M. K., Briker, L., Zhang, M., Baron, M., Heilmann, S., Deforet, M., Kenny, C., Ferretti, L. P., Huang, T.-H., Perlee, S., Garg, M., Nsengimana, J., Saini, M., Montal, E., Tagore, M., Newton- Bishop, J., … White, R. M. (2021). Cooperation between melanoma cell states promotes metastasis through heterotypic cluster formation. Developmental Cell, 56(20), 2808–2825.e10. 10.1016/j.devcel.2021.08.018

5. Chen, Y., Song, Y., Du, W., Gong, L., Chang, H., & Zou, Z. (2019). Tumor-associated macrophages: An accomplice in solid tumor progression. Journal of Biomedical Science, 26(1), 78. 10.1186/s12929-019-0568-z

6. Deng, Q., Yoo, S. K., Cavnar, P. J., Green, J. M., & Huttenlocher, A. (2011). Dual roles for Rac2 in neutrophil motility and active retention in zebrafish hematopoietic tissue. Developmental Cell, 21(4), 735–745. 10.1016/j.devcel.2011.07.013

7. Dillekås, H., Rogers, M. S., & Straume, O. (2019). Are 90% of deaths from cancer caused by metastases? Cancer Medicine, 8(12), 5574–5576. 10.1002/cam4.2474

8. Ellett, F., Pase, L., Hayman, J. W., Andrianopoulos, A., & Lieschke, G. J. (2011). Mpeg1 promoter transgenes direct macrophage-lineage expression in zebrafish. Blood, 117(4), e49–e56. 10.1182/blood-2010-10-314120

9. Giese, M. A., Hind, L. E., & Huttenlocher, A. (2019). Neutrophil plasticity in the tumor microenvironment. Blood, 133(20), 2159–2167. 10.1182/blood-2018-11-844548

10. Gonzalez, H., Hagerling, C., & Werb, Z. (2018). Roles of the immune system in cancer: From tumor initiation to metastatic progression. Genes & Development, 32(19– 20), 1267–1284. 10.1101/gad.314617.118

11. Goswami, S., Sahai, E., Wyckoff, J. B., Cammer, M., Cox, D., Pixley, F. J., Stanley, E. R., Segall, J. E., & Condeelis, J. S. (2005). Macrophages promote the invasion of breast carcinoma cells via a colony-stimulating factor-1/epidermal growth factor paracrine loop. Cancer Research, 65(12), 5278–5283. 10.1158/0008-5472.CAN-04-1853

12. Hafemeister, C., & Satija, R. (2019). Normalization and variance stabilization of single- cell RNA-seq data using regularized negative binomial regression. Genome Biology, 20(1), 296. 10.1186/s13059-019-1874-1

13. Hanna, S. J., McCoy-Simandle, K., Leung, E., Genna, A., Condeelis, J., & Cox, D. (2019). Tunneling nanotubes, a novel mode of tumor cell–macrophage communication in tumor cell invasion. Journal of Cell Science, 132(3), jcs223321. 10.1242/jcs.223321

14. Hao, Y., Stuart, T., Kowalski, M. H., Choudhary, S., Hoffman, P., Hartman, A., Srivastava, A., Molla, G., Madad, S., Fernandez-Granda, C., & Satija, R. (2024). Dictionary learning for integrative, multimodal and scalable single-cell analysis. Nature Biotechnology, 42(2), 293–304. 10.1038/s41587-023-01767-y

15. Heilmann, S., Ratnakumar, K., Langdon, E., Kansler, E., Kim, I., Campbell, N. R., Perry, E., McMahon, A., Kaufman, C., van Rooijen, E., Lee, W., Iacobuzio-Donahue, C., Hynes, R., Zon, L., Xavier, J., & White, R. (2015). A Quantitative System for Studying Metastasis Using Transparent Zebrafish. Cancer Research, 75(20), 4272–4282. 10.1158/0008-5472.CAN-14-3319

16. Hsu, A. P., Donkó, A., Arrington, M. E., Swamydas, M., Fink, D., Das, A., Escobedo, O., Bonagura, V., Szabolcs, P., Steinberg, H. N., Bergerson, J., Skoskiewicz, A., Makhija, M., Davis, J., Foruraghi, L., Palmer, C., Fuleihan, R. L., Church, J. A., Bhandoola, A., … Holland, S. M. (2019). Dominant activating RAC2 mutation with lymphopenia, immunodeficiency, and cytoskeletal defects. Blood, 133(18), 1977– 1988. 10.1182/blood-2018-11-886028

17. Huemer, K., Squirrell, J. M., Swader, R., LeBert, D. C., Huttenlocher, A., & Eliceiri, K. W. (2017). zWEDGI: Wounding and Entrapment Device for Imaging Live Zebrafish Larvae. Zebrafish, 14(1), 42–50. 10.1089/zeb.2016.1323

18. Hunter, M. V., Moncada, R., Weiss, J. M., Yanai, I., & White, R. M. (2021). Spatially resolved transcriptomics reveals the architecture of the tumor-microenvironment interface. Nature Communications, 12(1), 6278. 10.1038/s41467-021-26614-z

19. Joshi, S., Singh, A. R., Zulcic, M., Bao, L., Messer, K., Ideker, T., Dutkowski, J., & Durden, D. L. (2014). Rac2 controls tumor growth, metastasis and M1-M2 macrophage differentiation in vivo. PloS One, 9(4), e95893. 10.1371/journal.pone.0095893

20. Kitamura, T., Qian, B.-Z., & Pollard, J. W. (2015). Immune cell promotion of metastasis. Nature Reviews Immunology, 15(2), 73–86. 10.1038/nri3789

21. Kwan, K. M., Fujimoto, E., Grabher, C., Mangum, B. D., Hardy, M. E., Campbell, D. S., Parant, J. M., Yost, H. J., Kanki, J. P., & Chien, C.-B. (2007). The Tol2kit: A multisite gateway-based construction kit for Tol2 transposon transgenesis constructs. Developmental Dynamics : An Official Publication of the American Association of Anatomists, 236(11), 3088–3099. 10.1002/dvdy.21343

22. Li, L., Yan, B., Shi, Y.-Q., Zhang, W.-Q., & Wen, Z.-L. (2012). Live imaging reveals differing roles of macrophages and neutrophils during zebrafish tail fin regeneration. The Journal of Biological Chemistry, 287(30), 25353–25360. 10.1074/jbc.M112.349126

23. Linde, N., Casanova-Acebes, M., Sosa, M. S., Mortha, A., Rahman, A., Farias, E., Harper, K., Tardio, E., Reyes Torres, I., Jones, J., Condeelis, J., Merad, M., & Aguirre-Ghiso, J. A. (2018). Macrophages orchestrate breast cancer early dissemination and metastasis. Nature Communications, 9(1), 21. 10.1038/s41467-017-02481-5

24. Mishra, A. K., Rodriguez, M., Torres, A. Y., Smith, M., Rodriguez, A., Bond, A., Morrissey, M. A., & Montell, D. J. (2023). Hyperactive Rac stimulates cannibalism of living target cells and enhances CAR-M-mediated cancer cell killing. Proceedings of the National Academy of Sciences, 120(52), e2310221120. 10.1073/pnas.2310221120

25. Miskolci, V., Squirrell, J., Rindy, J., Vincent, W., Sauer, J. D., Gibson, A., Eliceiri, K. W., & Huttenlocher, A. (2019). Distinct inflammatory and wound healing responses to complex caudal fin injuries of larval zebrafish. eLife, 8. 10.7554/eLife.45976

26. Miskolci, V., Wu, B., Moshfegh, Y., Cox, D., & Hodgson, L. (2016). Optical Tools To Study the Isoform-Specific Roles of Small GTPases in Immune Cells. The Journal of Immunology, 196(8), 3479–3493. 10.4049/jimmunol.1501655

27. Paterson, N., & Lämmermann, T. (2022). Macrophage network dynamics depend on haptokinesis for optimal local surveillance. eLife, 11, e75354. 10.7554/eLife.75354

28. Roh-Johnson, M., Shah, A. N., Stonick, J. A., Poudel, K. R., Kargl, J., Yang, G. H., di Martino, J., Hernandez, R. E., Gast, C. E., Zarour, L. R., Antoku, S., Houghton, A. M., Bravo-Cordero, J. J., Wong, M. H., Condeelis, J., & Moens, C. B. (2017). Macrophage-Dependent Cytoplasmic Transfer during Melanoma Invasion In Vivo. Developmental Cell, 43(5), 549–562.e6. 10.1016/j.devcel.2017.11.003

29. Rosowski, E. E., Deng, Q., Keller, N. P., & Huttenlocher, A. (2016). Rac2 Functions in Both Neutrophils and Macrophages To Mediate Motility and Host Defense in Larval Zebrafish. The Journal of Immunology, 197(12), 4780–4790. 10.4049/jimmunol.1600928

30. Rosowski, E. E., Raffa, N., Knox, B. P., Golenberg, N., Keller, N. P., & Huttenlocher, A. (2018). Macrophages inhibit Aspergillus fumigatus germination and neutrophil- mediated fungal killing. PLoS Pathogens, 14(8), e1007229. 10.1371/journal.ppat.1007229

31. Roussos, E. T., Balsamo, M., Alford, S. K., Wyckoff, J. B., Gligorijevic, B., Wang, Y., Pozzuto, M., Stobezki, R., Goswami, S., Segall, J. E., Lauffenburger, D. A., Bresnick, A. R., Gertler, F. B., & Condeelis, J. S. (2011). Mena invasive (MenaINV) promotes multicellular streaming motility and transendothelial migration in a mouse model of breast cancer. Journal of Cell Science, 124(13), 2120–2131. 10.1242/jcs.086231

32. Schindelin, J., Arganda-Carreras, I., Frise, E., Kaynig, V., Longair, M., Pietzsch, T., Preibisch, S., Rueden, C., Saalfeld, S., Schmid, B., Tinevez, J.-Y., White, D. J., Hartenstein, V., Eliceiri, K., Tomancak, P., & Cardona, A. (2012). Fiji: An open- source platform for biological-image analysis. Nature Methods, 9(7), 676–682. 10.1038/nmeth.2019

33. Shin, M., Male, I., Beane, T. J., Villefranc, J. A., Kok, F. O., Zhu, L. J., & Lawson, N. D. (2016). Vegfc acts through ERK to induce sprouting and differentiation of trunk lymphatic progenitors. *Development (Cambridge*, England), 143(20), 3785–3795. 10.1242/dev.137901

34. Weber, M., Huisken, J., Schlaeppi, A., & Graves, A. (2021). Light Sheet Microscopy of Fast Cardiac Dynamics in Zebrafish Embryos. JoVE, 174, e62741. 10.3791/62741

35. Weiss, J. M., Hunter, M. V., Cruz, N. M., Baggiolini, A., Tagore, M., Ma, Y., Misale, S., Marasco, M., Simon-Vermot, T., Campbell, N. R., Newell, F., Wilmott, J. S., Johansson, P. A., Thompson, J. F., Long, G. V., Pearson, J. V., Mann, G. J., Scolyer, R. A., Waddell, N., … White, R. M. (2022). Anatomic position determines oncogenic specificity in melanoma. Nature, 604(7905), 354–361. 10.1038/s41586-022-04584-6

36. Wheeler, A. P., Wells, C. M., Smith, S. D., Vega, F. M., Henderson, R. B., Tybulewicz, V. L., & Ridley, A. J. (2006). Rac1 and Rac2 regulate macrophage morphology but are not essential for migration. Journal of Cell Science, 119(13), 2749–2757. 10.1242/jcs.03024

37. Wyckoff, J. B., Wang, Y., Lin, E. Y., Li, J., Goswami, S., Stanley, E. R., Segall, J. E., Pollard, J. W., & Condeelis, J. (2007). Direct Visualization of Macrophage- Assisted Tumor Cell Intravasation in Mammary Tumors. Cancer Research, 67(6), 2649–2656. 10.1158/0008-5472.CAN-06-1823

38. Xue, Q., Varady, S. R. S., Waddell, T. Q. A., Roman, M. R., Carrington, J., & Roh- Johnson, M. (2023). Lack of Paxillin phosphorylation promotes single-cell migration in vivo. The Journal of Cell Biology, 222(3). 10.1083/jcb.202206078

