## Supplementary figures and images for "Real-time imaging reveals a role for macrophage protrusive motility in melanoma invasion"

### Supplemental Figure 1

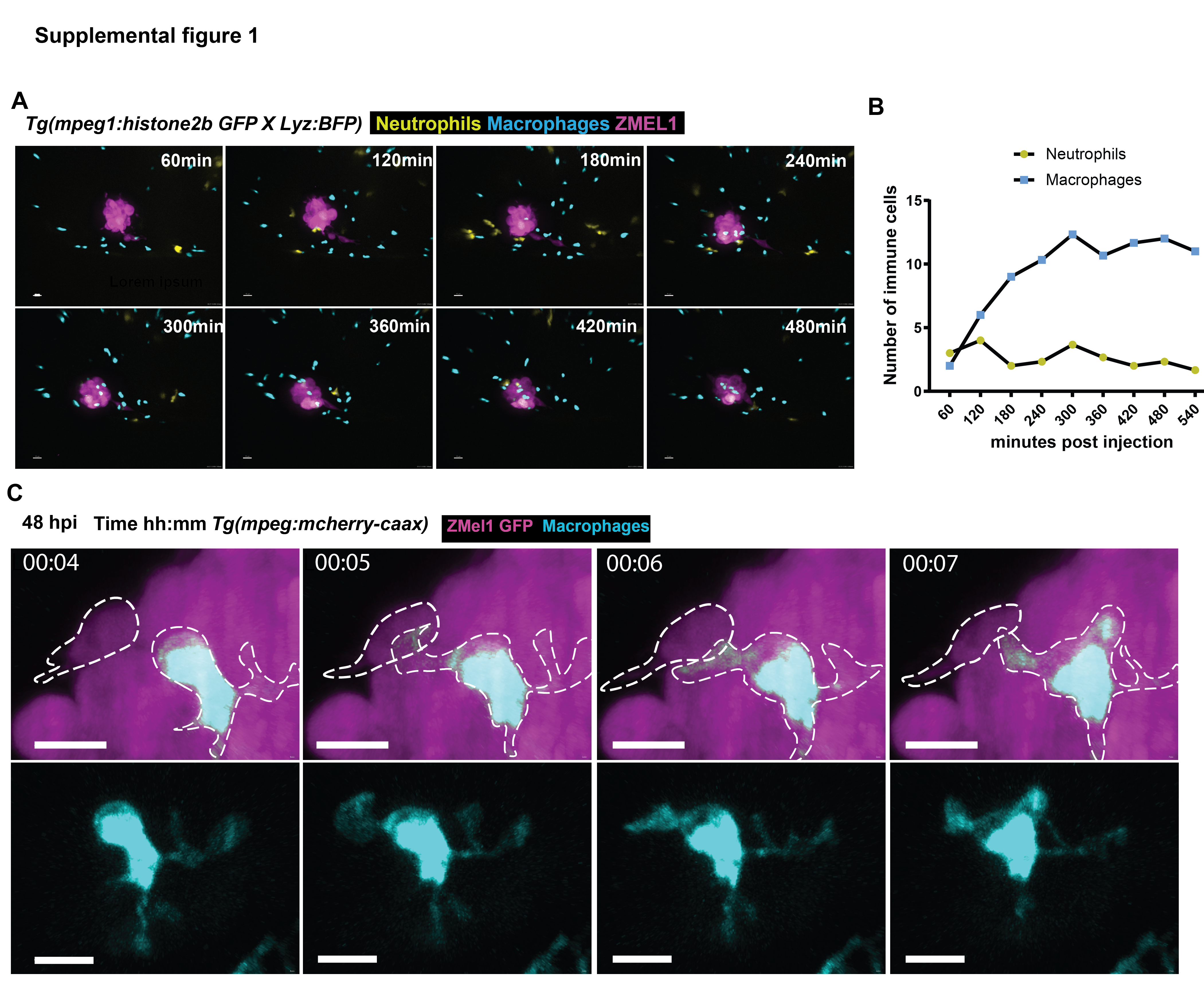

### Supplemental Figure 2

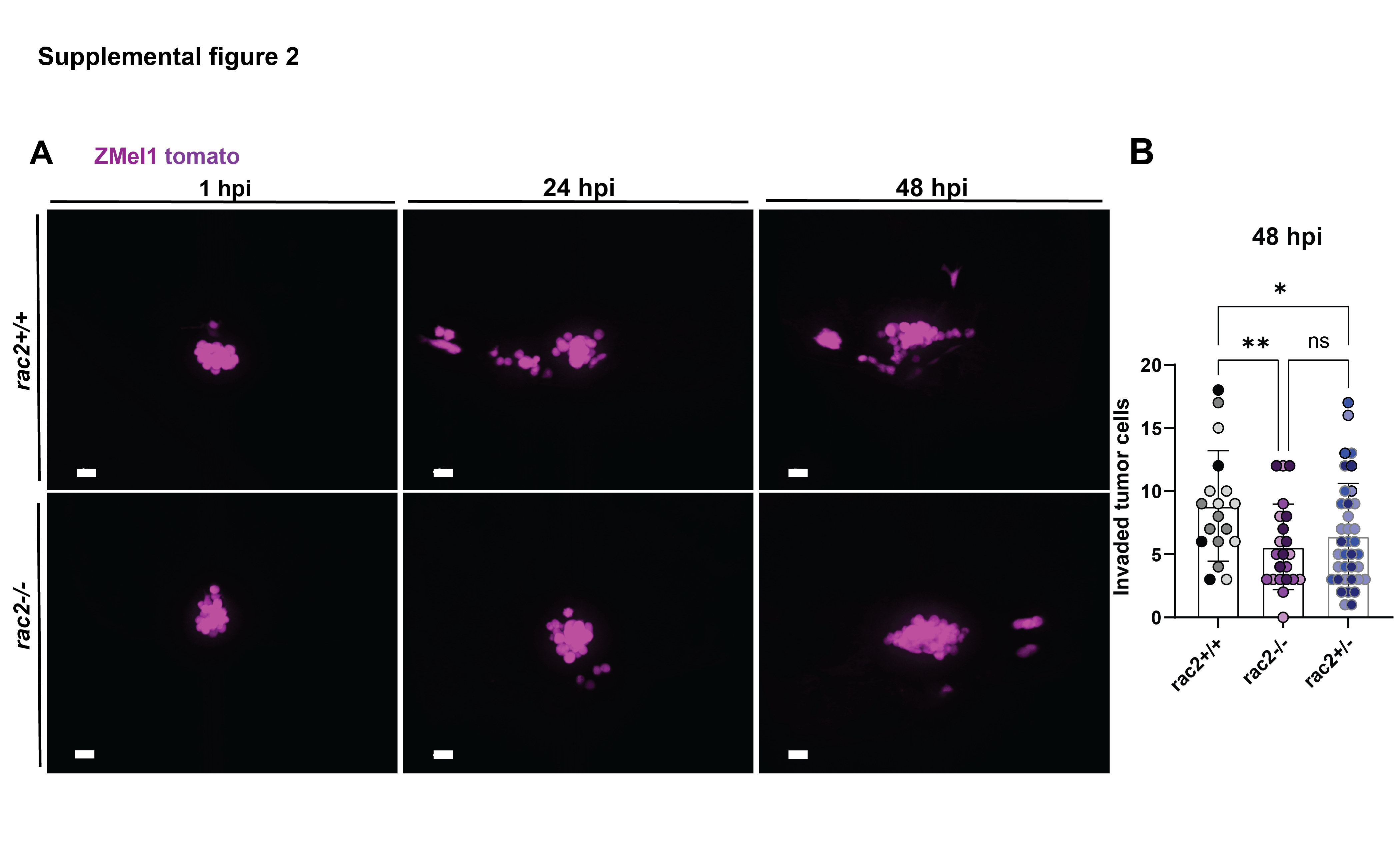
